# Variation in seagrass habitat use by fishery-important nekton across the Gulf of Mexico revealed by deep transfer learning with trophic group priors

**DOI:** 10.64898/2026.05.26.727508

**Authors:** Liying Li, Jon Rodemann, Christian Hayes, Benjamin Belgrad, Kelly M. Darnell, Charles W. Martin, Bradley T Furman, Delbert L. Smee, M. Zachary Darnell

## Abstract

Understanding how species use habitats across environmental gradients is central to guiding fisheries management and habitat restoration, yet inference is often limited by heterogeneous data and inconsistent observations. In coastal ecosystems, variation in relationships between nekton and seagrass habitats remains unresolved, in part because habitat structure, environmental context, and sampling methods are rarely integrated in predictive models. Here, we combine multi-gear monitoring data from across the Gulf of Mexico with a transfer-learning framework that incorporates trophic priors to quantify how nekton respond to seagrass structure under varying environmental conditions. We show that environmental gradients—including temperature, salinity, and water clarity—define broad-scale distributions, while seagrass structure refines habitat use at local scales. Apparent inconsistencies in seagrass–nekton relationships are largely attributable to environmental context and differences in observational processes. By integrating observations collected using different sampling methods, our approach reveals consistent species–environment relationships across sites and improves predictive performance, particularly for data-limited species, by using trophic priors. We further show that species differ in their responses to environmental gradients, with some exhibiting consistent patterns across sites and others showing strong context dependence. These results demonstrate that combining heterogeneous datasets can strengthen ecological inference and provide a pathway for scalable, data-driven conservation and restoration in rapidly changing coastal systems.

## 1. Introduction

Research on nekton habitat use in coastal ecosystems consistently highlights the critical role of structured habitats—particularly seagrass meadows, salt marshes, and mangroves—as nursery and foraging grounds (Vassallo et al., 2014). Seagrass habitats generally support higher densities, diversity, and production of juvenile fishes and mobile invertebrates than adjacent unvegetated substrates, reinforcing their importance for ecosystem functions that support wildlife (Whaley et al., 2023). However, these habitats are increasingly threatened by climate change and land-use pressures, including ocean warming and climate-driven environmental change (Waycott et al., 2009), sea-level rise (Capistrant-Fossa et al., 2024), eutrophication and declining water quality (Waycott et al., 2009; Thomsen et al., 2020), and coastal development (Waycott et al., 2009), all of which have contributed to widespread degradation and loss of seagrass ecosystems. Despite growing recognition of seagrass habitat importance and its restoration needs, effective restoration actions remain limited by insufficient information to support informed decision-making, particularly in understanding how habitat structure interacts with environmental context to shape nekton species habitat use across different environments (Sheaves et al., 2024; Preston et al., 2025; Sievers et al., 2024).

There is extensive research on nekton use of seagrass habitats, demonstrating that within-patch structural attributes—such as shoot density, canopy height, and above– and belowground biomass—are consistently linked to nekton abundance and community composition (Orth et al., 1984a,b; Heck et al., 1995; Gray et al., 1996, French et al., 2021). Numerous studies have also shown that nekton habitat use is shaped by adjacent habitats and environmental conditions, including interactions between seagrass and marsh edges, dissolved oxygen constraints, and broader habitat mosaics that influence recruitment and community composition (Gilpin, 2023; Clark et al., 2025; Endresz et al., 2026). However, studies that extend beyond within-patch structure to landscape-scale patch attributes have produced weak or inconsistent results, often due to species– and time-dependent variability, scale mismatches, and methodological limitations such as sampling artifacts and survey design (Underwood et al., 2000). In addition, recent work highlights that differences in sampling gear and observational methods can interact with habitat heterogeneity to shape observed nekton patterns, further complicating inference (Wang et al., 2025). Insufficient comparisons across environmental gradients using a consistent sampling protocol have led to uncertainty about whether the observed variability in Nekton habitat use arises from spatial variation in seagrass features or from differences in environmental context, sampling design, and scale (Sheaves et al., 2024). As a result, the relative importance of seagrass structure versus broader environmental drivers remains unresolved across systems.

Sampling design has long been recognized as a major source of uncertainty in detecting ecological patterns (Underwood et al., 2000; Wiens, 1989; Levin, 1992; Thrush et al., 1997). Recent coordinated data collection efforts, such as those funded by the NOAA RESTORE program, apply consistent sampling schemes across environmentally distinct locations, reducing methodological variability and enabling more reliable cross-site comparisons. However, different sampling gears are often deployed to cover varying water depths and habitats. These gears exhibits selective efficiency across taxa, sizes, and microhabitats, thereby biasing observed community structure and obscuring underlying ecological relationships (Underwood et al., 2000; Bradley et al., 2012).

These biases are further amplified in heterogeneous environments, where habitat complexity interacts with sampling methods, making it difficult to distinguish the ecological signals from observation processes (Moua et al., 2020; Wang et al., 2025). In addition, ecological datasets are often characterized by imbalanced species observations, limiting the ability to robustly model responses across entire assemblages. Traditional approaches, such as random forest models, typically require larger and more balanced datasets and tend to fit the local data distribution more closely, particularly when data are sparse (Pichler and Hartig, 2023).

Here, we address this gap by quantifying how nekton respond to seagrass structural features that vary across large-scale environmental gradients spanning multiple sites in the Gulf of Mexico using deep learning approaches. Rather than focusing on patch configuration, we explicitly control for co-varying environmental drivers (e.g., hydrological conditions, surrounding habitat context, and climatic variation) and evaluate how relationships between seagrass attributes (including shoot structure and biomass components) and species occurrence change across sites. Deep learning with shared representations enables the integration of multi-gear species observation data and the learning of complex, nonlinear relationships between habitat structure and species occurrence, rather than site-specific artifacts, improving cross-site inference (LeCun et al., 2015; Reichstein et al., 2019). Transfer learning helps the model focus on generalizable ecological patterns under sparse and heterogeneous observations (Pan & Yang, 2010; Weiss et al., 2016; Ge et al., 2023; He and Garcia, 2014). However, these techniques have been rarely applied to empirical fish surveys, needless to say, nekton and seagrass interactions.

We leverage recent advances in deep learning and developed a transfer learning framework (Pan & Yang, 2010), in which models were first trained on higher-sample trophic groups to learn general habitat–ecological relationships and then adapted to species-level prediction (Weiss et al., 2016). This approach allows species-level inference to build upon hierarchical representations that capture interactions among environmental variables and species communities, which are then transferred to species-level habitat selection (Olden et al., 2008; Willcock et al., 2018).

The combination of standardized sampling and integrative modeling allows us to move beyond site-specific approaches and directly evaluate ecological relationships across systems. Our objective is to quantify how nekton responses to seagrass habitat structure vary across environmental gradients and to disentangle the relative roles of within-patch structure and broader environmental context. More broadly, this framework provides a pathway to improve predictive understanding of habitat–species relationships in data-limited, heterogeneous monitoring environments, enabling more reliable cross-site comparisons of nekton habitat use. These advances are critical for informing conservation and restoration strategies, as they allow managers to identify context-dependent seagrass habitat functions, prioritize areas for restoration, and better anticipate fish responses under ongoing environmental change.

## 2. Methods

### 2.1. Study area and data sampling

Seagrass ecosystems provide critical habitat for nekton, offering food resources, shelter, and foraging grounds for a wide range of species. This study investigated seagrass habitat use by eight fishery-important species across six regions of the Gulf of Mexico to evaluate spatial and temporal variation in habitat selection. The six study sites included Lower Laguna Madre (LM), Coastal Bend (CB), Chandeleur Islands (LA), St. George Sound (SG), Cedar Key (CK), and Charlotte Harbor (CH). At each site, approximately 20–25 sampling stations were established using a tessellated hexagon sampling design to ensure spatially balanced coverage.

Sampling was conducted across multiple seasonal periods. Two baseline sampling seasons occurred in 2018 (May 14–June 14 and August 13–September 17). Following disturbance from Hurricane Michael, three sites (CK, SG, and LA) were additionally sampled in October 2018 (post-hurricane), May–June 2019, and August–September 2019 to capture post-disturbance habitat dynamics.

Nekton were sampled using two complementary methods targeting different components of the water column. Trawl sampling targeted mid– and upper-water nekton using 2–3 minute tows conducted at speeds of 3.5–5.5 km h ¹. Organisms were sorted to species, counted, weighed, and, for up to 25 individuals per species, measured. Epibenthic sled sampling targeted bottom-dwelling nekton. The sled was deployed at the midpoint of each trawl and manually towed over a standardized distance of 13.3 m. Samples were returned to the laboratory for species identification, enumeration, and measurement.

Seagrass complexity metrics were measured in association with both sampling methods. For trawl-associated sampling, seagrass percent cover by species and shoot density were recorded using four quadrats at each of the three points along each trawl path. Canopy height was measured on three representative shoots per quadrat. For sled-associated sampling, seagrass cores were collected at three points along the sled path. Core samples were analyzed to quantify blade length, blade width, total leaf area, leaf area index (LAI), aboveground biomass, belowground biomass, and epiphyte biomass. Detailed descriptions of all nekton and seagrass methodology can be found in Hayes (2021).

Nekton and seagrass datasets were integrated at the haul level using unique sampling identifiers (pull_id and tow_id). The final curated dataset comprised 645 haul-level observations, including 322 epibenthic sled hauls and 323 trawl hauls. Each unique haul was treated as a single modeling unit. Species abundance values were converted to occurrence, which was defined as binary presence (1) when species abundance is non-zero, and absence (0) otherwise.

The focal predicted species comprised eight fishery management-relevant fisheries species: Brown shrimp (*Penaeus aztecus*), Pink shrimp (*Penaeus duorarum*), White shrimp (*Litopenaeus setiferus*), Blue crab (*Callinectes sapidus*), Red drum (*Sciaenops ocellatus*), Spotted seatrout (*Cynoscion nebulosus*), Gray snapper (*Lutjanus griseus*), and Lane snapper (*Lutjanus synagris*).

### 2.2. Predictors and missing values

Predictor variables were collected along both sled and trawl sampling. In addition to seagrass habitat structure, we constructed 4 categories of predictors to represent environmental conditions, spatial context, and observation processes. All predictors were aggregated to the haul level to match the response variables. Predictor categories include environmental variables: light attenuation, depth, Secchi depth, temperature, salinity, dissolved oxygen, and macroalgae biomass (macro_weight); Spatial coordinates: start and end latitude/longitude of each haul; **S**ampling gear indicator (gear_is_sled); temporal variables: year, month, day-of-year (derived from sampling date); categorical context variables: one-hot encoded site, station, sampling period, and substrate categories. We compared models with and without gear_is_sled as a predictor to decide how to make the model make full use of mixed gear data.

Missing data are pervasive in long-term ecological monitoring due to evolving protocols, variable sampling effort, and logistical constraints in field and laboratory measurements. Rather than removing incomplete observations or predictors, we adopted a missingness-aware modeling approach to retain all available ecological information.

For each numeric predictor, missing values were handled using a two-part representation. First, median imputation was applied to fill missing numeric values. Second, a corresponding missingness indicator variable was created (1 = missing, 0 = observed). This approach allows the model to distinguish between true observed values and imputed values, thereby preserving information about data availability and potential observation bias. All numeric predictors were standardized after imputation.

For categorical predictors, missing values were not imputed numerically; instead, they were assigned to an explicit “Missing” category prior to one-hot encoding. Temporal variables were derived from raw date fields (e.g., year, month, day-of-year). When date parsing failed, the resulting missing values were handled using the same numeric imputation and indicator strategy described above. For missing scientific names, corresponding common names were used where available. If both identifiers were missing, the record was assigned to an “Unknown species” category.

**Table 1.** Number of predictors in each category after filling in the missing data.

| Predictor block | Raw features | Missingness indicators | Total columns used |
| --- | --- | --- | --- |
| Environmental variables | 7 | 7 | 14 |
| Spatial coordinates | 4 | 4 | 8 |
| Gear indicator: gear_is_sled | 1 | 1 | 2 |
| Temporal variables | 3 | 3 | 6 |
| Seagrass structure variables | 165 | 165 | 330 |
| Categorical context variables | 151 | 0 | 151 |
| Total | 331 | 180 | 511 |

### 2.3. Two-stage hierarchical modeling framework

There are eight focal species: Brown shrimp (*Penaeus aztecus*), Pink shrimp (*P. duorarum*), White shrimp (*L. setiferus*), Blue crab (*Callinectes sapidus*), Red drum (*Sciaenops ocellatus*), Spotted seatrout (*Cynoscion nebulosus*), Gray snapper (*Lutjanus griseus*), and Lane snapper (*L. synagris*). These focal species exhibited highly imbalanced sampling frequencies, with occupancy ranging from 5 (Red drum) to over 200 (Brown shrimp), reflecting substantial ecological and observational heterogeneity (Table 2). To address this, we developed a two-stage hierarchical modeling framework to characterize habitat selection by fishery-important nekton while accounting for class imbalance and heterogeneity in observations.

**Table 2.** The sampling occupancy of each of the targeted fishery-important nekton species.

| Common name | Scientific name | Sampling occupancy |
| --- | --- | --- |
| Brown shrimp | *Penaeus aztecus* | 206 |
| Pink shrimp | *Penaeus duorarum* | 187 |
| White shrimp | *Litopenaeus setiferus* | 10 |
| Blue crab | *Callinectes sapidus* | 151 |
| Red drum | *Sciaenops ocellatus* | 5 |
| Spotted seatrout | *Cynoscion nebulosus* | 83 |
| Gray snapper | *Lutjanus griseus* | 60 |
| Lane snapper | *Lutjanus synagris* | 21 |

In the first stage, the model was pretrained to predict the occurrence of 11 trophic groups using environmental and seagrass predictors, which serve as trophic-level priors for subsequent species-level prediction and habitat selection (Table 3). Compared to individual species, trophic groups have substantially higher sample sizes, enabling the model to learn stable relationships between habitat conditions and ecological function. All nekton species other than the eight focal species were assigned to these trophic groups based on feeding type and ecological position (e.g., detritivore demersal, piscivore demersal, planktivore pelagic), using published literature and ecological databases.

**Table 3.** The sampling occupancy of each of the trophic groups.

| Group | Explanation | Occupancy |
| --- | --- | --- |
| Omnivore demersal | Bottom-associated omnivores that feed on a mixed diet of animal and non-animal material. | 565 |
| Detritivore demersal | Bottom-associated consumers that feed mainly on detritus or organic material in or on sediments. | 457 |
| Carnivore demersal | Bottom-associated animal-feeding consumers that prey broadly on other animals. | 378 |
| Carnivore epifaunal | Animal-feeding consumers living on or immediately above the substrate or vegetation surface. | 331 |
| Detritivore epifaunal | Surface-associated consumers that feed mainly on detritus or organic material on vegetation or substrate surfaces. | 317 |
| Piscivore demersal | Bottom-associated fish-eating consumers. | 126 |
| Planktivore epifaunal | Surface-associated consumers feed mainly on plankton. | 95 |
| Invertivore demersal | Bottom-associated consumers feed mainly on invertebrates. | 65 |
| Carnivore pelagic | Water-column animal-feeding consumers that prey broadly on other animals. | 52 |
| Planktivore pelagic | Water-column consumers feed mainly on plankton. | 49 |
| Piscivore pelagic | Water-column fish-eating consumers. | 41 |

This stage trains a shared encoder that captures general ecological structure supporting major functional groups, effectively learning a habitat template. The pretrained encoder is subsequently fine-tuned to predict the occurrence of the eight focal fisheries species. Because the pretraining and fine-tuning tasks differ in output dimensionality (11 trophic groups vs. 8 species), only the shared encoder and latent layers are transferred, while the species-specific output head is newly initialized. The model is then optimized on species-level occurrence data. This transfer-learning strategy allows species predictions to build upon biologically informed representations rather than learning from sparse data alone.

### 2.4. Hierarchical ecological consistency constraint

To preserve the ecological structure learned during pretraining, we introduce a hierarchical ecological consistency loss that links species predictions to their parent trophic groups. Each focal species is mapped to a corresponding trophic group (e.g., shrimp and crab to detritivore demersal; red drum to omnivore demersal; snapper and seatrout to piscivore demersal). During fine-tuning, the pretrained trophic-group model is retained as a frozen teacher, and an auxiliary loss penalizes discrepancies between species-level predictions and the teacher’s group-level outputs.

This constraint enforces consistency between fine-scale species predictions and broader functional guild structure. As a result, species-level habitat selection is learned as a combination of shared habitat associations inherited from the trophic group and species-specific deviations from that ecological template. This hierarchical regularization improves stability under data sparsity and enhances the ecological interpretability of the hierarchical model with trophic-level priors.

### 2.5. GMVAE with contrastive learning (sub-model)

The underlying predictive model is a supervised, label-conditioned Gaussian Mixture Variational Autoencoder (GMVAE) with contrastive learning (Figure 1), following Junwen Bai et al. (2020). The model maps environmental and seagrass predictors into a latent Gaussian space using an MLP feature encoder that outputs mean and variance parameters. Label (trophic groups or species) –dependent structure is incorporated through a label encoder that defines label-specific embeddings as a Gaussian mixture prior. For multilabel samples, the latent Gaussian mixture prior is constructed as a weighted combination of active label components.

**Figure 1.**
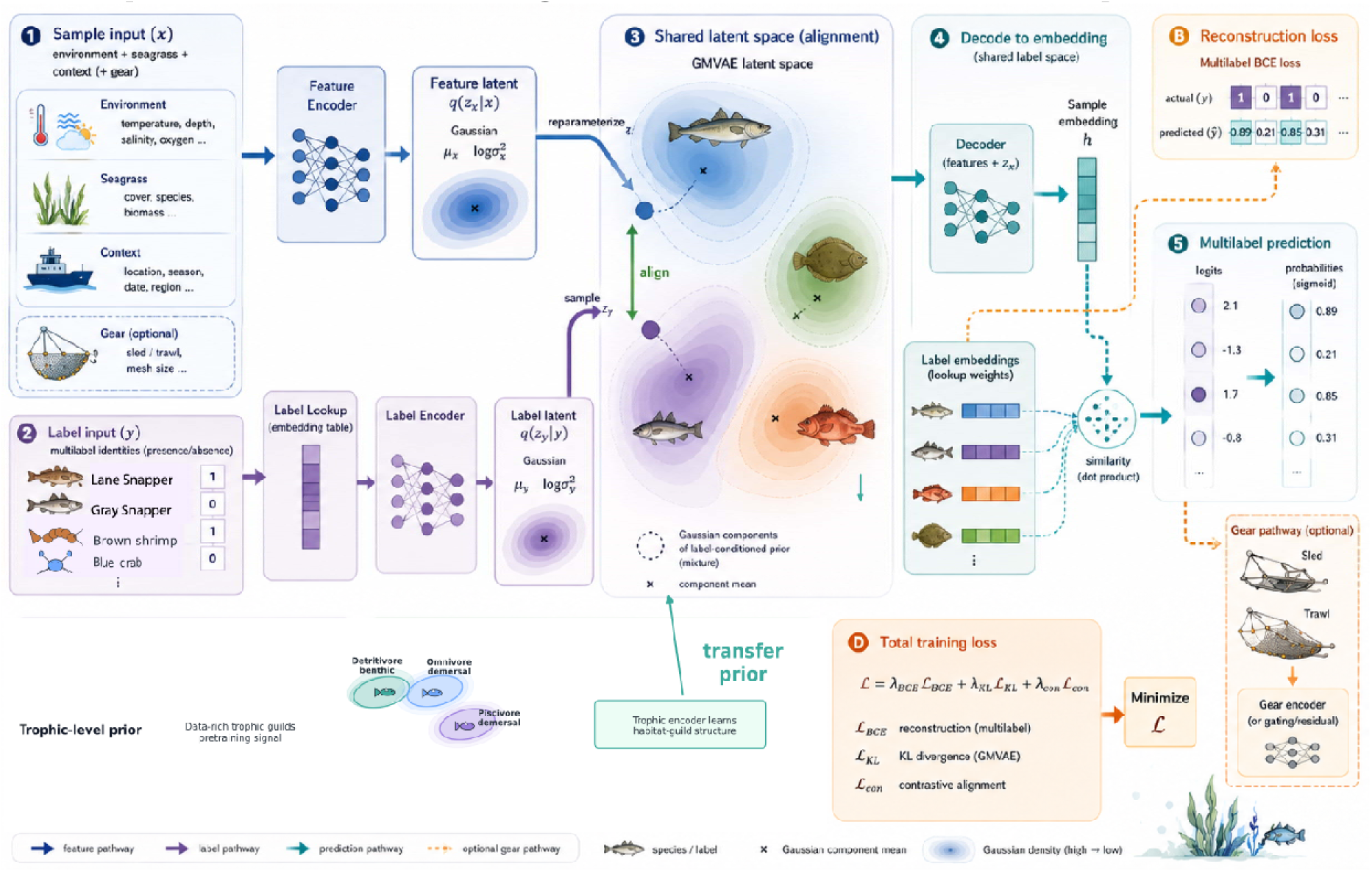
GMVAE with the transfer learning of the trophic level priors method diagram.

Data were partitioned into training, validation, and test sets using a fixed random seed (SEED = 1), with approximate proportions of 72%, 18%, and 10%, respectively. To address class imbalance, we employed class-balanced minibatch sampling with rare-positive oversampling to ensure adequate representation of infrequent species during training.

Model training optimizes three objectives: (i) KL divergence, (ii) multilabel classification loss (binary cross-entropy), and (iii) contrastive loss, which aligns feature embeddings with label embeddings and separates ecologically distinct classes. The contrastive objective improves representation quality by encouraging samples sharing similar ecological characteristics to cluster in the latent space, while separating dissimilar samples.

### 2.6. Model evaluation

Model performance was assessed using complementary threshold-independent and threshold-dependent metrics, including AUPR, ROC AUC, and F1, as well as multilabel metrics (micro– and macro-F1). During validation, predictions were evaluated across a grid of decision thresholds (0.1–0.9), and for each metric, the optimal threshold was retained. Model selection was based on validation macro-F1, such that reported validation performance reflects threshold-optimized rather than fixed-threshold predictions.

Testing followed the same procedure, reporting the best values across thresholds for Hamming accuracy, example-based F1, micro-F1, macro-F1, and precision at 1. This approach emphasizes performance under optimal decision calibration while avoiding reliance on an arbitrary threshold. We compared the performance between models, whether using gear types as a model feature (predictor), and chose the best model for analyzing Nekton seagrass habitat use variations.

### 2.7. Habitat selection analysis

We used gradient-based attribution to infer species–environment relationships from model predictions. For each species and sample, integrated gradients were computed with respect to the input environmental features, yielding local attribution scores that quantify the contributions of predictors. Because attributions are defined at the sample level, they can be aggregated spatially to reveal regional patterns. We averaged attributions within regions and compared their relative magnitudes across space, allowing us to identify shifts in dominant environmental drivers. This approach provides a flexible, model-based analog to habitat selection analysis, enabling direct comparison of how species–environment associations vary across heterogeneous landscapes.

### 2.8. Partial dependence analysis

Partial dependence analysis was based on the fitted no-gear transfer-learning model and the predictor importance analysis. For top environmental predictors, including salinity, dissolved oxygen, temperature, seagrass richness, and secchi depth, each variable was varied independently over a grid spanning its 5th–95th percentile original range, while all other predictors were held at their observed values for each sample. Model predictions were recomputed across the grid and averaged over samples to produce species-specific mean predicted probability response curves. Predictor values were back-transformed to original field units before generating partial dependency plots (PDP) to facilitate ecological interpretation.

For top (importance) seagrass within-patch features, including shoots, above and below biomass, canopy, leaf width, and leaves per shoot, PDPs were constructed at the level of functional features (e.g., shoots, canopy, above– and belowground biomass, leaf width, and leaves per shoot) by aggregating these variables across seven different seagrass species where species were present. Feature-level values (mean) were varied over their 5th–95th percentile range and redistributed across constituent predictors while preserving within-feature composition. Model predictions across the evaluated gradient were then computed and averaged across samples to obtain mean probability prediction response curves. It is worthwhile to note all PDPs represent changes in mean predicted probability rather than thresholded occurrence.

## 3. Results

### 3.1. Model performance

Model performance varied across species, reflecting differences in sampling support and class imbalance (Table 4). Overall, the model achieved strong predictive performance for well-sampled species, including Pink shrimp (*Penaeus duorarum*; ROC-AUC = 0.91, F1 = 0.79), Spotted seatrout (*Cynoscion nebulosus*; ROC-AUC = 0.88, F1 = 0.73), Brown shrimp (*P. aztecus*; ROC-AUC = 0.85, F1 = 0.69), and Blue crab (*Callinectes sapidus*; ROC-AUC = 0.84, F1 = 0.71). Gray snapper (*Lutjanus griseus*) also showed strong discrimination ability (ROC-AUC = 0.89), although F1 scores were moderately lower (0.67), reflecting class imbalance.

**Table 4.** Species-level model performance metrics with the optimized thresholds.

| Species | Scientific name | Support | roc_auc | aupr | f1 |
| --- | --- | --- | --- | --- | --- |
| Brown shrimp | *Penaeus aztecus* | 23 | 0.85 | 0.74 | 0.69 |
| Pink shrimp | *Penaeus duorarum* | 24 | 0.91 | 0.83 | 0.79 |
| White shrimp | *Litopenaeus setiferus* | 1 | 0.811 | 0.08 | 0 |
| Blue crab | *Callinectes sapidus* | 22 | 0.84 | 0.67 | 0.71 |
| Red drum | *Sciaenops ocellatus* | 2 | 0.71 | 0.53 | 0.5 |
| Spotted seatrout | *Cynoscion nebulosus* | 11 | 0.88 | 0.82 | 0.73 |
| Gray snapper | *Lutjanus griseus* | 8 | 0.89 | 0.82 | 0.67 |
| Lane snapper | *Lutjanus synagris* | 1 | 0.98 | 0.5 | 0.67 |

In contrast, rare species exhibited unstable or reduced performance, particularly White shrimp (*L. setiferus*) and Red drum (*Sciaenops ocellatus*), which had extremely low support and correspondingly low or inconsistent F1 and AUPR values. These patterns highlight the challenges of modeling species with limited observations despite the use of transfer learning from trophic group prior.

Across all evaluation metrics, the mixed-gear model outperformed the model that included gear as a simple feature, particularly under the per-gear dynamic threshold (Table 5). The per-gear dynamic threshold refers to selecting an optimal classification threshold separately for each sampling gear, rather than applying a single fixed threshold (e.g., 0.5) across all data and species (Supplementary Information Figure S1). The mixed model achieved higher overall performance (HA = 0.92, ebF1 = 0.76, miF1 = 0.80, p@1 = 0.80) compared to the gear-as-feature model (HA = 0.88, ebF1 = 0.67, miF1 = 0.72, p@1 = 0.71). Improvements were especially evident in F1-based metrics and ranking performance (p@1), indicating better discrimination and prioritization of species occurrence probabilities.

**Table 5.** Test metrics comparison between models with and without the Gear type as a feature.

| Model | Threshold | ACC | HA | ebF1 | miF1 | maF1 | p@1 |
| --- | --- | --- | --- | --- | --- | --- | --- |
| Gear_is_Sled as a feature | per-gear dynamic | 0.4000 | 0.8808 | 0.6697 | 0.7232 | 0.5989 | 0.7077 |
| Gear_is_Sled as a feature | Fixed 0.5 | 0.3385 | 0.8865 | 0.5746 | 0.7122 | 0.5939 | 0.5846 |
| Gear mixed model | per-gear dynamic | 0.5385 | 0.9154 | 0.7565 | 0.7982 | 0.6206 | 0.8000 |
| Gear mixed model | Fixed 0.5 | 0.2769 | 0.8788 | 0.4535 | 0.6519 | 0.4245 | 0.5385 |

Although both models enabled the sled prediction to borrow the prediction strength from trawl data, which is a more abundant dataset, the gear mixed model avoided using gear type as a prediction shortcut, amplifying the importance of other predictors. Thus, this gear mixed model produced the best balanced performance across gear types especially at the optimized thresholds (Table 6). Under a fixed threshold (0.5), performance declined for both models, but the mixed model remained more sensitive to threshold choice, with larger reductions in F1 metrics (Table 5), suggesting that dynamic thresholding is important for capturing class imbalance and gear-specific detection patterns.

**Table 6.**
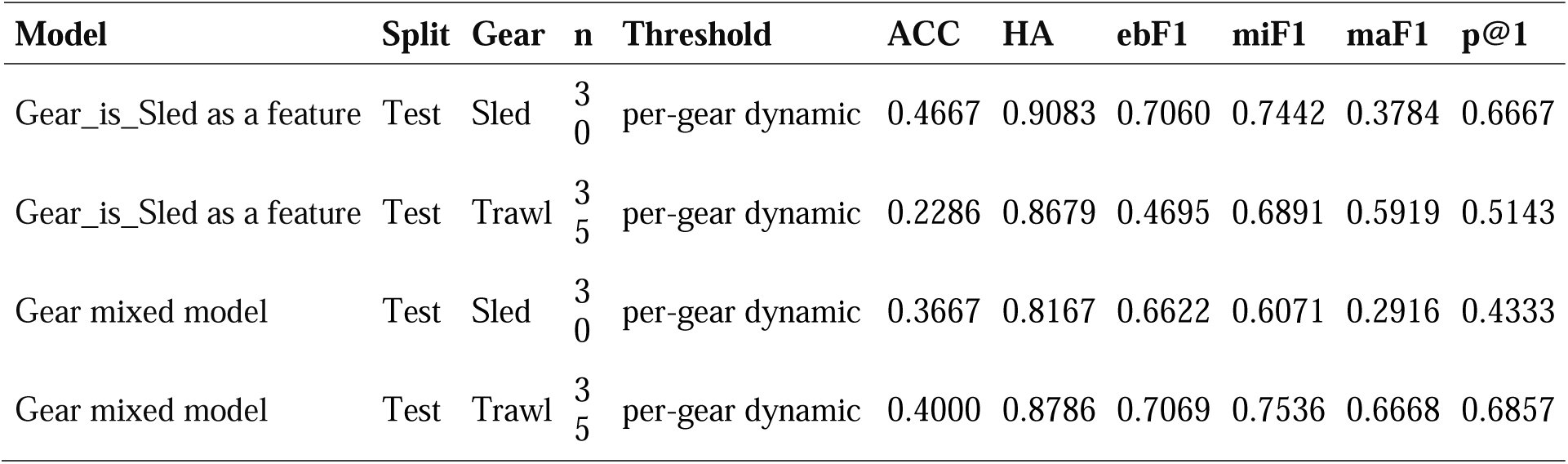
Test metrics split by gear type comparison between models with and without the Gear type as a feature, using a per-gear dynamic threshold.

At the species level, performance varied between sled and trawl data, but the mixed model showed improved or comparable performance across gears, particularly for trawl observations (Supplementary Information Table S1). Both models failed to predict white shrimp and red drum who have extremely low sample support in either scenarios. We, therefore, removed these shrimp and red drum from the target species list for the subsequent analysis.

### 3.2. Predictor importance

Feature attribution analysis revealed both consistent environmental drivers and differences in signal structure between species-level and trophic-level models (Figure 2). Across species, the most important predictors were seagrass richness, Secchi depth, temperature, dissolved oxygen, and macroalgal biomass, highlighting the role of environmental gradients in shaping distributions. Strong species-specific sensitivities were evident (e.g., temperature for Blue crab (0.26) and DO for Pink shrimp (0.22)), whereas rarer species showed weaker signals. At the trophic-group level, predictor importance distributions were broader and more diffuse, reflecting aggregation across species with diverse ecological roles. Nevertheless, the same core environmental drivers include salinity, temperature, water clarity, seagrass richness, and dissolved oxygen, remaining consistently important across groups. Peak predictor importance values were lower than species-level values but still highlighted key ecological relationships, for example, DO remained consistently important, particularly for demersal groups.

**Figure 2.**
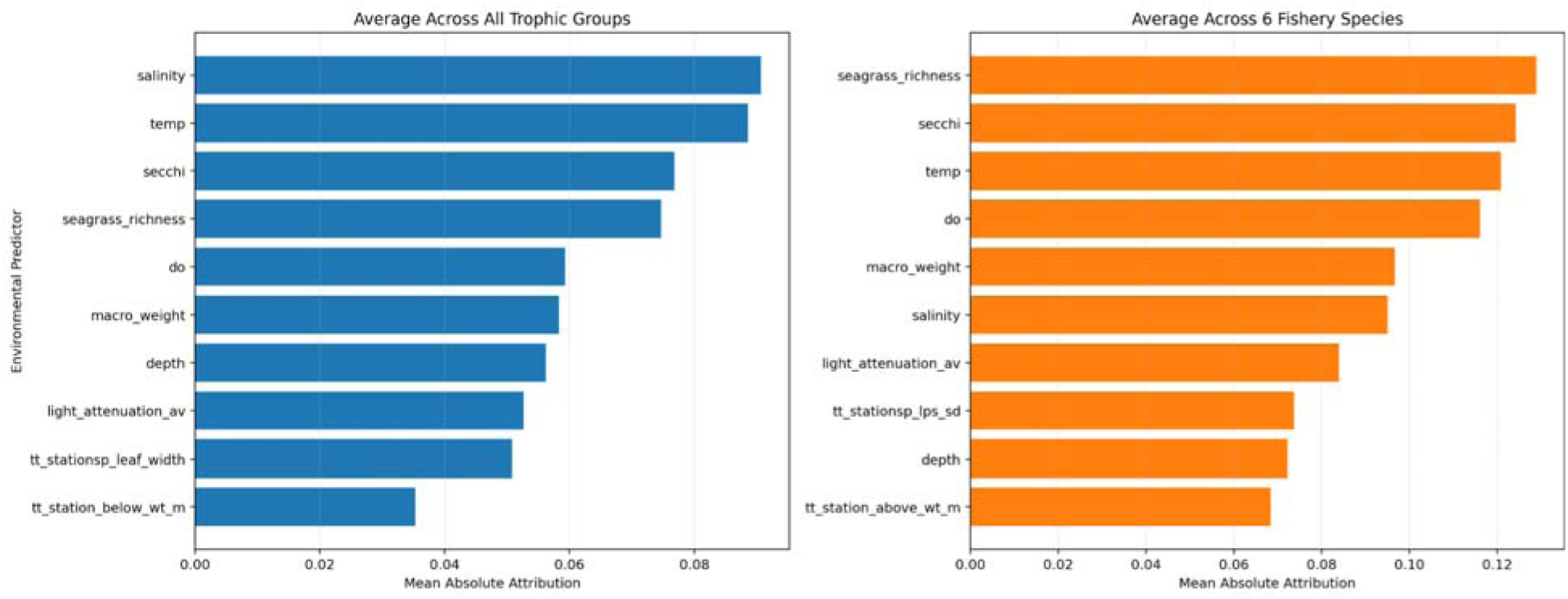
Environment and seagrass predictors importance barplots to 11 trophic groups and 6 target nekton species across all six sites.

### 3.3. Site-level variation in seagrass habitat importance

Seagrass species (Figure 3) and feature importance (Figure 4) varied across sites, but several consistent patterns emerged in the role of seagrass species composition and structural attributes. Across all sites, *Thalassia testudinum* (TT) was consistently important for all species, highlighting its dominant role in shaping habitat use. *Syringodium filiforme* (SF) was also important at multiple sites (e.g., AP, CK), while *Halodule wrightii* (HW) contributed primarily to invertebrate responses (Figure 3).

**Figure 3.**
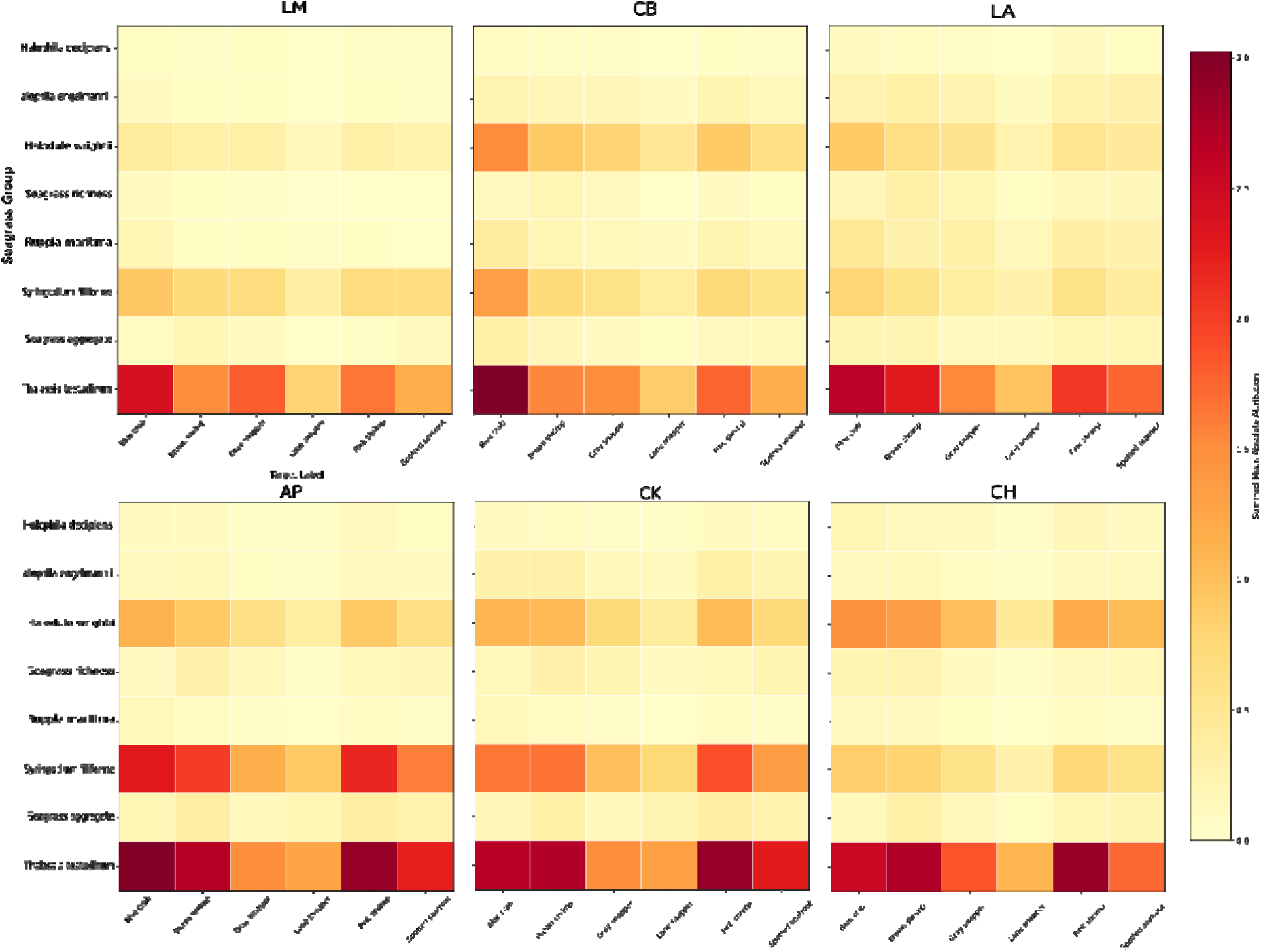
Seagrass species importance heatmap for 6 fishery-important species at all six sites – In AP and CK, Syringodium is also driving these species presences, while it is only Thalassia at the other 4 sites.

**Figure 4.**
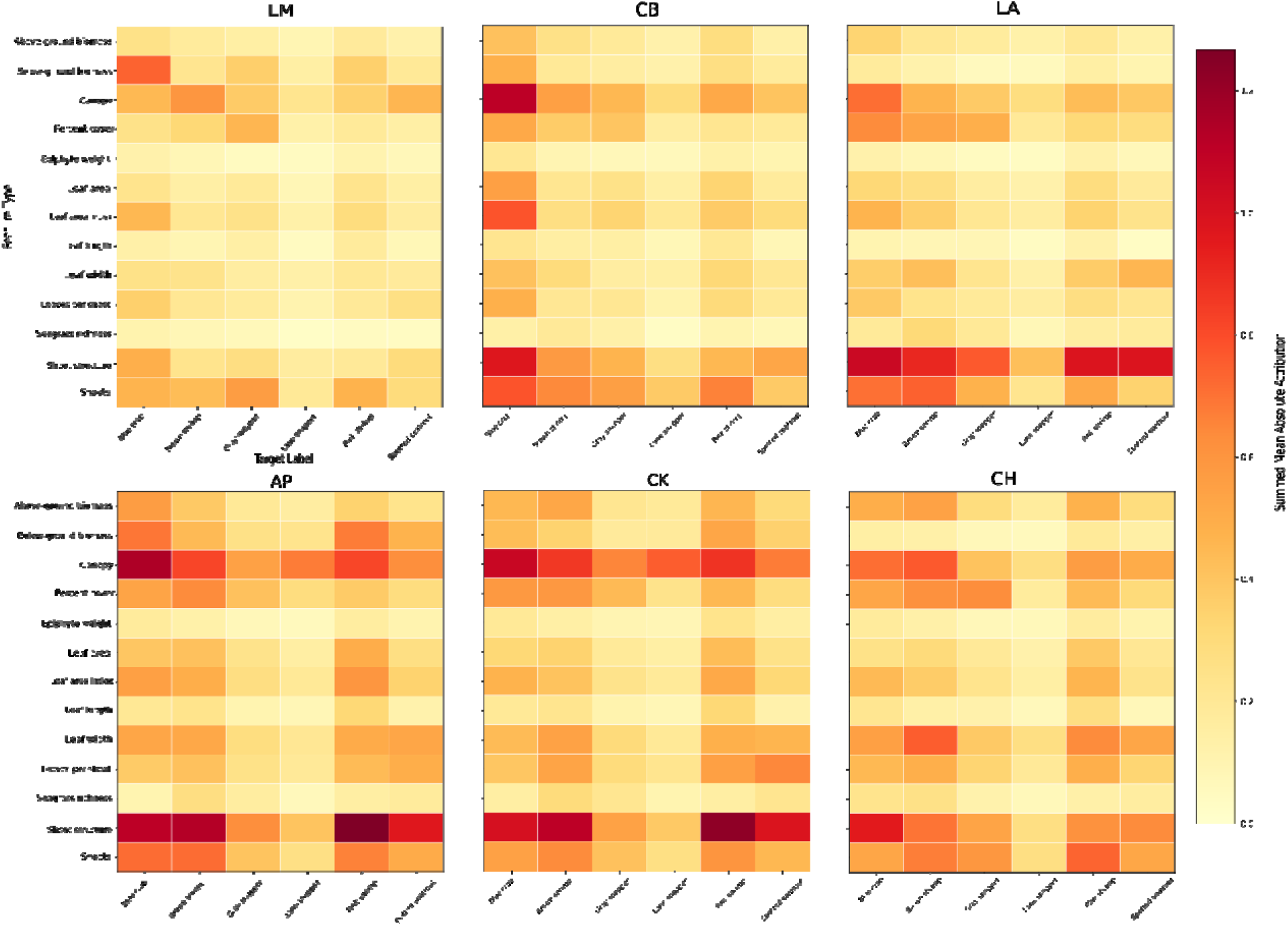
Seagrass habitat structure charactoristics importance heatmap for 6 fishery-important species at all six sites.

At Apalachicola (AP), both TT and SF were important across species, with shoot density, canopy, and shoot structure emerging as key predictors, alongside depth and macroalgal biomass for invertebrates (Figure 4). In Coastal Bend (CB), environmental drivers such as temperature and depth were important for blue crab, and DO for pink shrimp, while structural features (canopy, shoots, shoot structure) were most relevant for crabs, reflecting a community dominated by fewer taxa.

In Charlotte Harbor (CH), structural complexity (canopy, shoot structure, leaf width) was important across species, while environmental variables (depth, salinity, macroalgal biomass) were particularly influential for invertebrates. Cedar Key (CK) showed strong contributions from both environmental (DO, temperature, Secchi, richness) and structural variables, with canopy and shoot structure consistently important across taxa (Figure 4).

In Louisiana (LA), shoot density and shoot structure were dominant predictors across species, with canopy and percent cover contributing strongly to crabs and some fish species. In contrast, Lower Laguna Madre (LM) was distinguished by the importance of water clarity (Secchi), salinity, and DO, with canopy and belowground biomass important for some taxa, but notably little influence of shoot structure, unlike other sites.

### 3.4. Habitat use variation across sites between shrimp species

Partial dependence profiles revealed both consistent and site-specific responses of Brown shrimp (*Penaeus aztecus*) to environmental and seagrass predictors across the study region. Salinity showed a consistent negative relationship across all sites, indicating a reduced occurrence at higher salinity (Figure 5). Dissolved oxygen (DO) was generally positively associated with occurrence, except in the Texas sites (Lower Laguna Madre, LM; Coastal Bend, CB), where responses diverged, showing a negative relationship in LM and a nonlinear (parabolic) response in CB. Temperature exhibited predominantly positive effects across four of six sites, though responses were nonlinear in some cases, with a negative relationship observed in Louisiana (LA) and a parabolic response in LM. Among water clarity and habitat indicators, Secchi depth and seagrass richness were consistently positively associated with Brown shrimp occurrence across all sites, suggesting a preference for clearer waters and more diverse seagrass assemblages.

**Figure 5.**
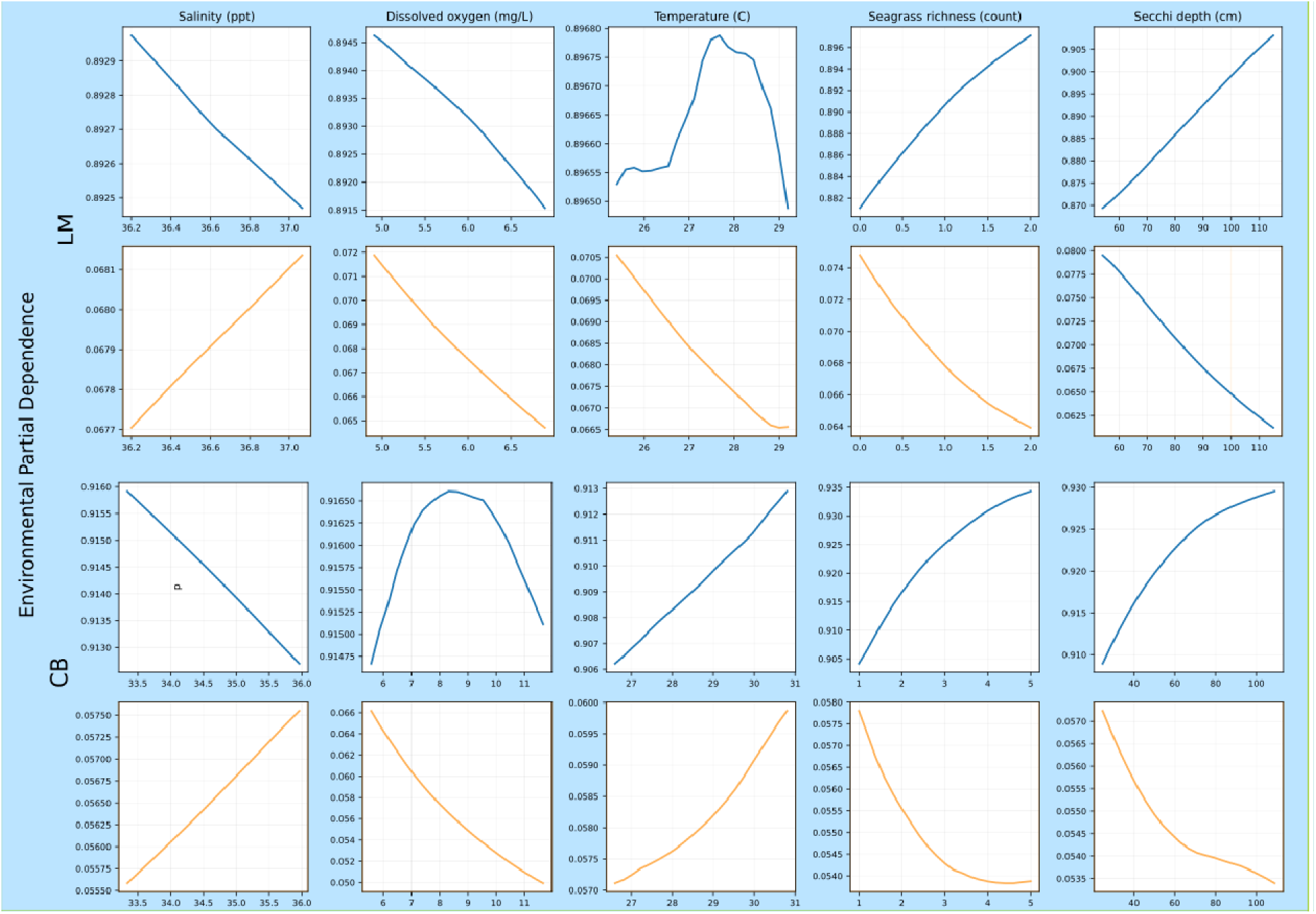
Environmental variable partial dependency in LM and CB for brown and pink shrimp.

Environmental variable partial dependence profiles for Pink shrimp (*Penaeus duorarum*) revealed patterns that were largely opposite to those observed for Brown shrimp, particularly for environmental drivers. Salinity showed a consistent positive relationship across all sites, while dissolved oxygen (DO) was negatively associated with occurrence throughout the study region (Figure 5). Temperature responses were more variable, with positive relationships in Coastal Bend (CB), Apalachicola (AP), and Louisiana (LA), and negative relationships in Lower Laguna Madre (LM), Cedar Key (CK), and Charlotte Harbor (CH), indicating strong spatial heterogeneity in thermal responses. Water clarity and habitat context also contrasted with Brown shrimp patterns. Both seagrass richness and Secchi depth were consistently negatively associated with Pink shrimp occurrence across sites, suggesting a preference for less diverse or more turbid environments relative to Brown shrimp.

Responses to seagrass structural attributes further emphasized contrasting habitat use by brown shrimp and pink shrimp, where structural seagrass attributes showed more complex patterns. Shoot density was generally negatively associated with brown shrimp occurrence, indicating reduced use of denser vegetation (Figure 6). Although shoot density was generally negatively associated with pink shrimp occurrence, in CH and CB, this pattern was weaker or reversed to a positive association for pink shrimp. Both brown and pink shrimp showed a positive relationship with aboveground biomass at most sites, except at CB. Belowground biomass exhibited high variability across sites, with no consistent pattern for either species.

**Figure 6.**
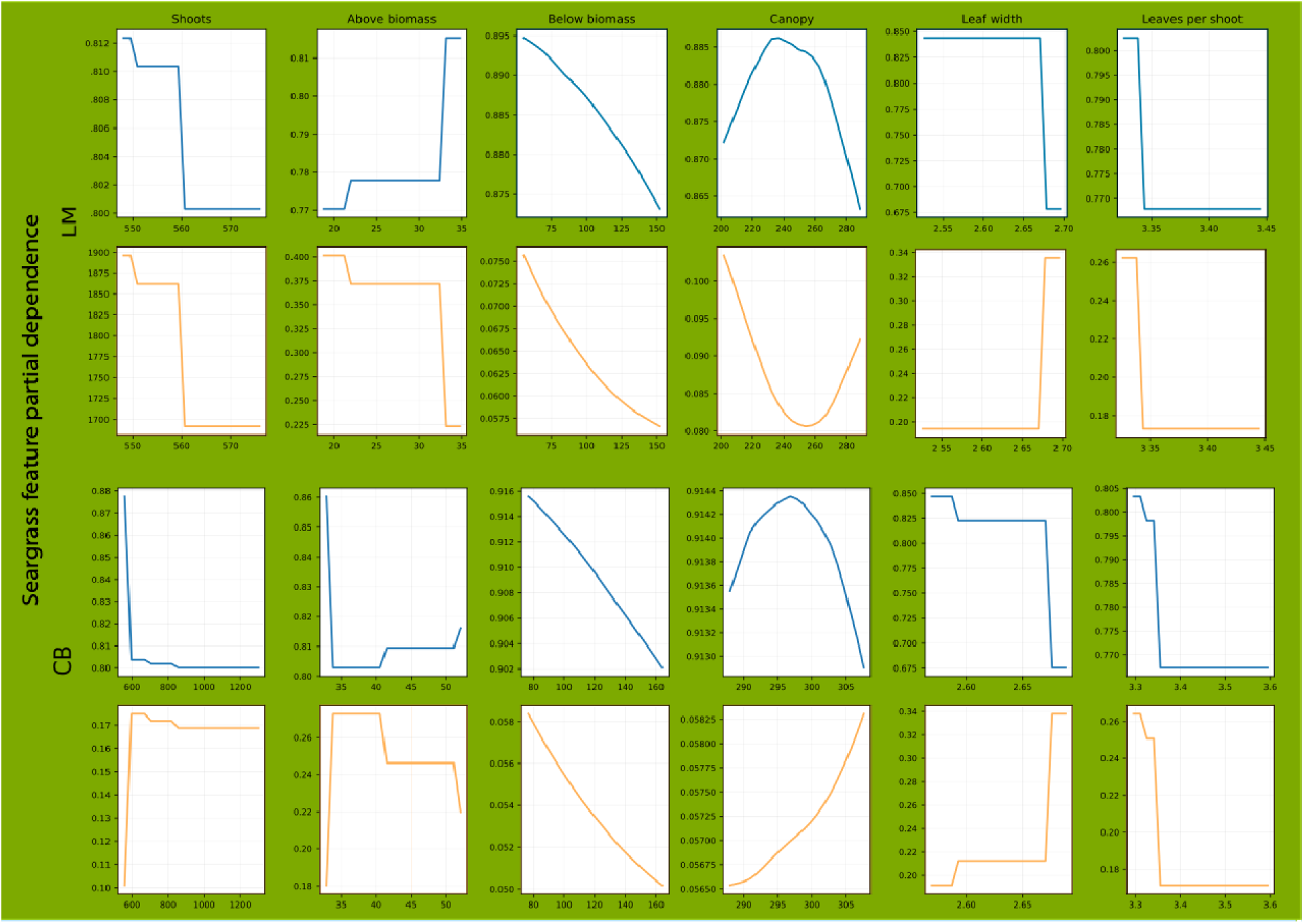
Seagrass habitat structure partial dependency in LM and CB for brown and Pink shrinp.

Canopy height and leaf width were generally negatively associated with Brown shrimp occurrence, except in Texas sites where responses were nonlinear (parabolic). In contrast to Brown shrimp, canopy height and leaf width were generally positively associated with pink shrimp occurrence, except in CH (negative) and LM (nonlinear, negative parabolic), indicating increased Pink shrimp occurrence in areas with broader and taller leaves in most sites.

Overall, Pink shrimp exhibit systematic differences in habitat associations relative to Brown shrimp, with most environmental and structural relationships reversed between species. Notably, temperature responses were the most variable across sites, suggesting strong spatial gradients in thermal conditions across the Gulf. These contrasting patterns provide evidence of niche differentiation between the two shrimp species, where species-specific responses to environmental gradients and habitat structure may reduce overlap and support coexistence across heterogeneous coastal systems.

### 3.5. Similar responses to environmental conditions and structural seagrass attributes by the blue crab and the Spotted seatrout

Blue crab (*Callinectes sapidus*) and Spotted seatrout (*Cynoscion nebulosus*) showed strong consistency in their responses to environmental variables across sites (Figure 7). Blue crab responses are closely aligned with those observed for Pink shrimp. Salinity exhibited a consistent positive relationship with Blue crab, but was generally negatively associated with Spotted seatrout occurrence.

**Figure 7.**
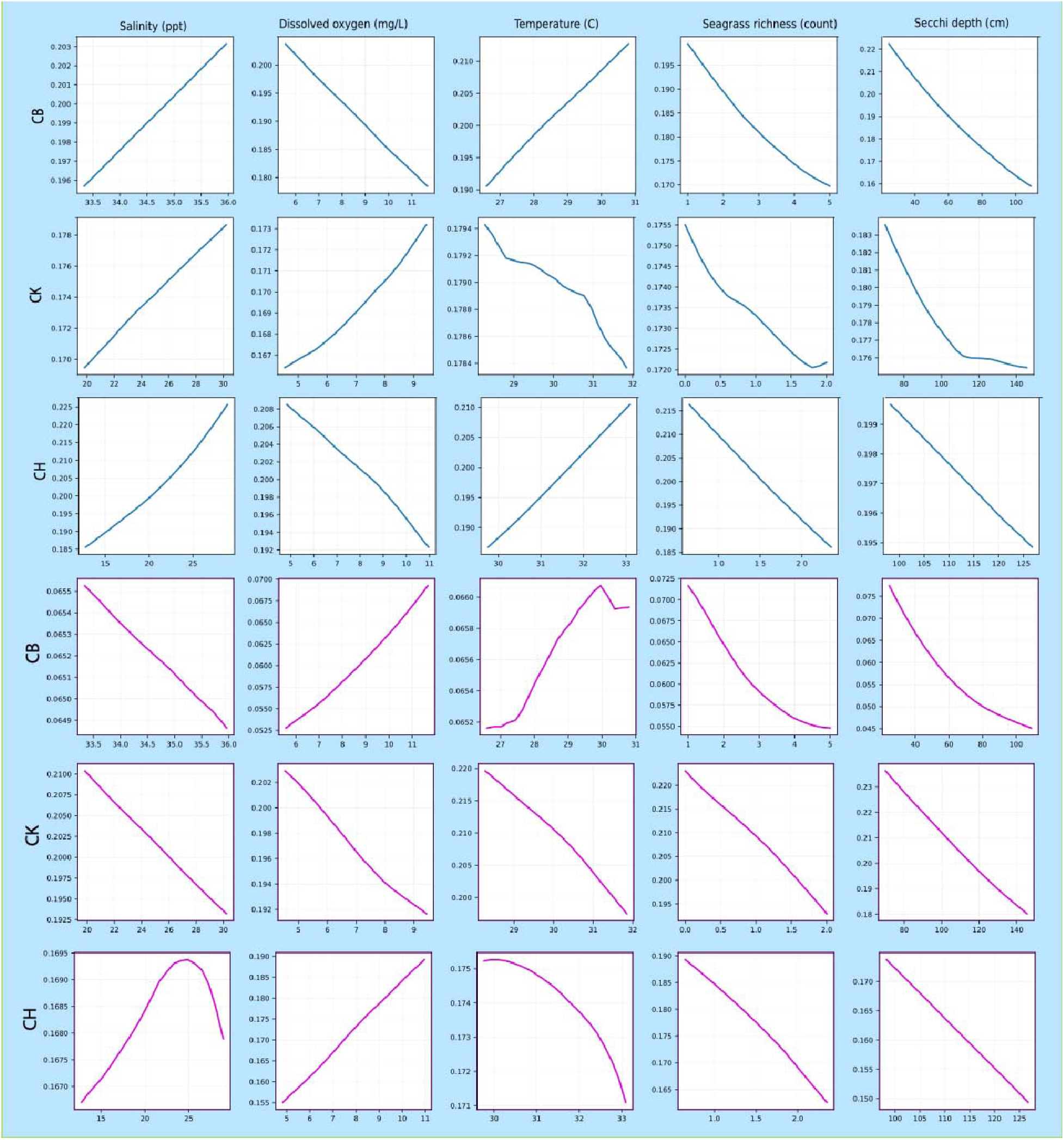
Blue crab (Blue Line) and Spotted Seatrout (Pink Line) environmental predictors, partial dependency across all sites.

Dissolved oxygen (DO) was generally negatively associated with both Blue crab and Spotted seatrout occurrence across all sites, with the exception of Cedar Key (CK) showing a positive association for Blue crab. Temperature was positively correlated with Blue crab across most sites, with CK again showing a weaker or contrasting response. Spotted seatrout has a clear threshold response to temperature, with occurrence increasing up to approximately 30°C (≈28°C in CK) and declining beyond this point, producing a consistent unimodal (parabolic) relationship across sites. Both blue crab and Spotted Seatrout have negative responses to increasing seagrass richness and Secchi depth across all sites, indicating a preference for more turbid conditions and less diverse seagrass assemblages.

Structural seagrass attributes showed stronger but more nuanced patterns across sites for Blue Crab and Spotted Sea Trout (Figure 8). Their structural seagrass attribute responses shared closely resembling patterns. Leaf width was consistently positively associated with the occurrence probability of both species across sites. Shoot density was generally positively associated with the occurrence of both species, suggesting a preference by both Blue crab and Seatrout for areas with higher shoot cover. In contrast, aboveground biomass was typically negatively associated with both species across all sites, except for seatrout in Coastal Bend (CB), indicating selection for habitats with structural elements (e.g., shoots) but lower overall biomass. Canopy height exhibited mixed responses. Both species have positive relationships with Canopy height in Louisiana (LA), CK, and Apalachicola (AP). In CH, Spotted seatrout shows negative relationships, whereas Blue crab shows a nonlinear response. In CB, the Blue crab shows a negative relationship, whereas the Spotted seatrout shows a nonlinear response. Overall, both species prefer fresher, warmer waters, while Spotted seatrout has an optimal temperature near 30°C. Both species select habitats with high shoot density but relatively low biomass, suggesting that they may favor structurally accessible habitats that balance cover and mobility. Site-specific interactions between canopy structure and environmental conditions determine the habitat suitability.

**Figure 8.**
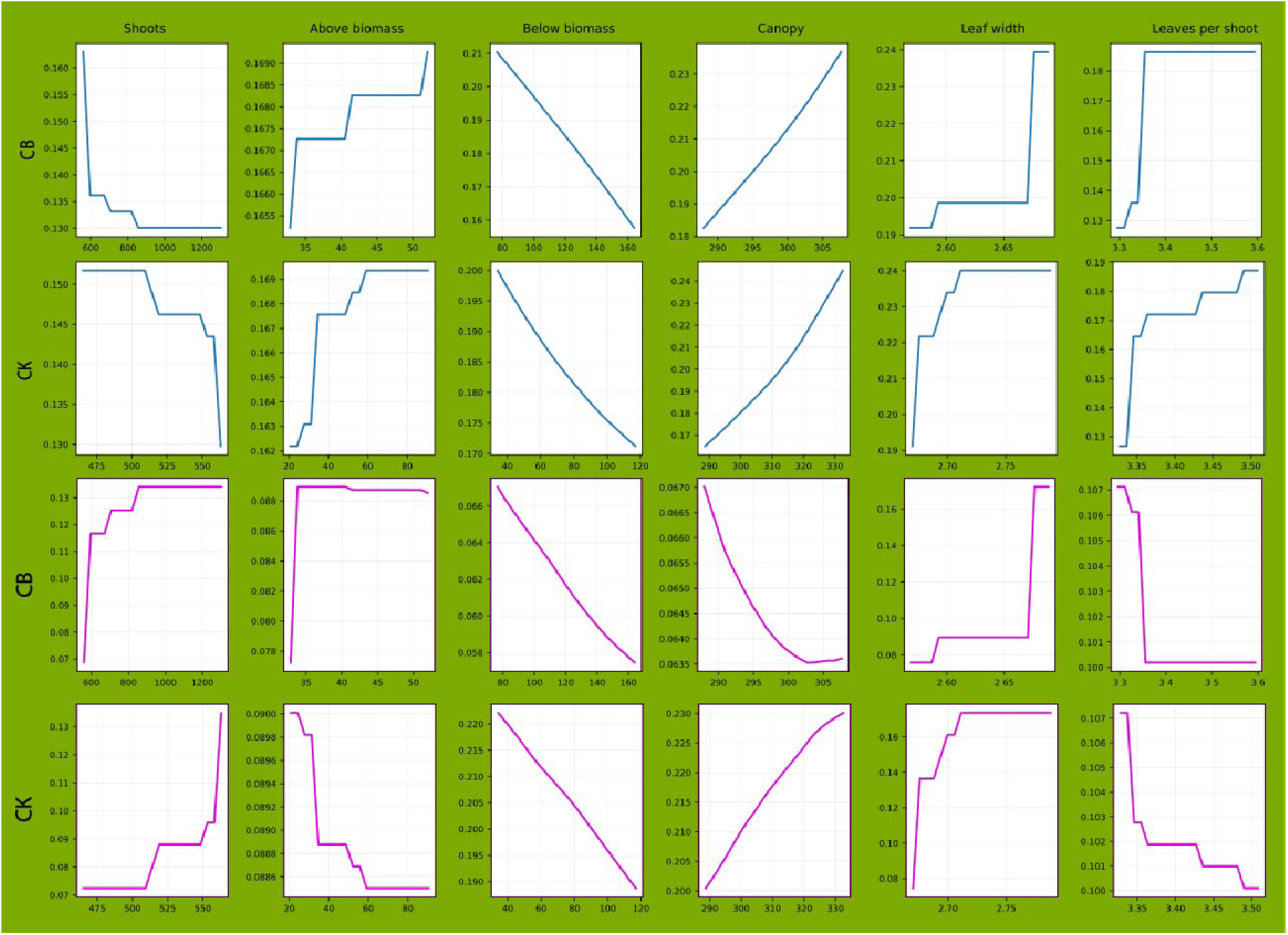
Blue crab (Blue Line) and Spotted Seatrout (Pink Line) structural seagrass feature partial dependency across in CK and CB.

### 3.6. Variable environmental condition response by Gray snapper across sites

Gray snapper (Lutjanus griseus), however, showed strong spatial variability in responses to environmental conditions across sites (Figure 9). Salinity was generally positively associated with occurrence, except in Cedar Key (CK), where the relationship was negative. Dissolved oxygen (DO) exhibited highly variable responses, with positive associations in Florida sites (Apalachicola, AP; CK; Charlotte Harbor, CH), negative relationships in Coastal Bend (CB), and nonlinear (parabolic) responses in Louisiana (LA) and Lower Laguna Madre (LM). Temperature responses were similarly heterogeneous, with positive relationships in AP and LM, negative relationships in CK, CH, and CB, and a nonlinear response in LA. Both seagrass richness and Secchi depth were negatively associated with occurrence across all sites, suggesting a preference for more turbid and less diverse seagrass environments. Although Gray snapper exhibits high context dependence in environmental responses, it has relatively consistent selection for specific components of seagrass structure, suggesting that structural habitat features may play a more stable role than environmental conditions in shaping habitat use by Gray snapper.

**Figure 9.**
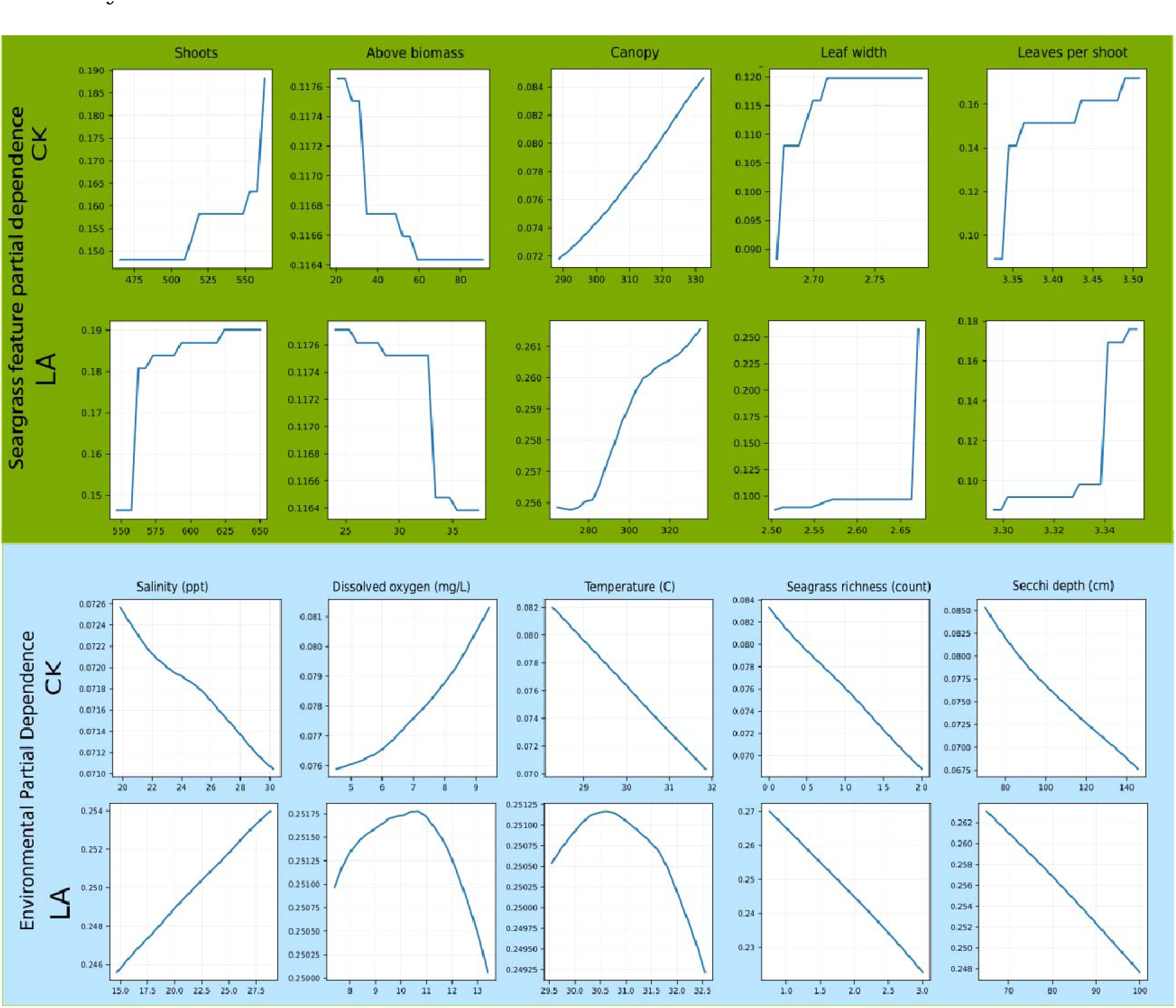
Seagrass within-patch feature and environmental context partial dependency in CK and LA for Gray Snapper.

## 4. Discussion

This study addresses a central challenge in coastal ecology: how to integrate heterogeneous observations and environmental context to understand species–habitat relationships across systems. By combining standardized sampling, multi-gear data integration, and a deep transfer learning framework, we provide new insight into how nekton respond to seagrass habitat structure across environmental gradients.

### 4.1. Resolving inconsistent seagrass–nekton relationships

Previous studies have reported inconsistent relationships between seagrass structure and nekton habitat use, often attributing variability to weak ecological effects or methodological limitations (Orth et al., 1984; Moccia et al., 2021; Gray et al., 1996; Frech et al., 2021). Our results suggest a different interpretation that these inconsistencies largely arise from unaccounted environmental heterogeneity and context dependence, consistent with ecological theory on scale and pattern (Wiens, 1989; Levin, 1992) and aligned with recent seascape perspectives emphasizing environmental gradients and connectivity (Sheaves et al., 2024; Preston et al., 2025).

Structural attributes such as shoot density, canopy, and biomass consistently influenced habitat use, but their effects varied depending on environmental context. This aligns with recent work showing that habitat complexity interacts with environmental stressors and landscape context to shape coastal communities (Unsworth et al., 2015; Sievers et al., 2024). Together, these findings indicate that seagrass structure operates within, rather than independently of, broader environmental gradients.

### 4.2. Species-specific responses and niche differentiation

Species-level analyses revealed clear evidence of niche differentiation, particularly among shrimp species, where Brown shrimp and Pink shrimp exhibited largely opposing responses to key environmental gradients. Such partitioning is consistent with recent empirical work demonstrating that species segregate along environmental gradients in coastal systems (Sheaves et al., 2024).

In contrast, Blue crab showed highly consistent responses across sites, with similar directional relationships for both environmental and structural predictors, suggesting a generalist strategy with broad tolerance for environmental conditions. A consistent affinity for structurally complex habitats was observed for Clue crab. Fish species exhibited mixed patterns, with Spotted seatrout showing a clear thermal optimum and Gray snapper exhibiting higher context dependence. These differences highlight that species vary not only in seagrass habitat preference but also in sensitivity to environmental context. Such context has been proven to be a key driver of community assembly in dynamic coastal systems (Sievers et al., 2024).

### 4.3. Environmental gradients dominate seagrass structure refines habitat use

Feature attribution results revealed a **hierarchical structure of drivers**, where environmental variables—temperature, salinity, water clarity, dissolved oxygen, and seagrass richness—dominate across both species and trophic levels. This is consistent with recent syntheses showing that environmental conditions strongly regulate coastal ecosystem functioning and species distributions (Waycott et al., 2009; zu Ermgassen et al., 2020; Sheaves et al., 2024).

Within this environmental envelope, seagrass structure refines habitat selection at local scales, supporting findings that structural complexity enhances refuge and foraging opportunities but operates within broader environmental constraints (Heck et al., 2003; Unsworth et al., 2015).

### 4.4. Advantages of integrating heterogeneous data with deep learning

The mixed-gear model outperformed models that treated gear as a simple covariate, supporting recent work emphasizing the importance of explicitly modeling observation processes (Isaac et al., 2014; Moua et al., 2020; Pichler et al., 2023) and avoiding model shortcuts in which gear effects are implicitly absorbed into ecological predictors. When gear is included only as a feature, the model may conflate differences in detection with true ecological signals, leading to biased inference and reduced generalization across sampling contexts.

Deep learning enables this integration by learning hierarchical representations that separate shared ecological structure from gear-specific observation patterns. In this framework, lower-level representations capture broad environmental gradients (e.g., temperature, salinity, habitat context), while higher-level representations encode interactions among habitat structure, species responses, and observation processes. As a result, patterns learned from different gears are aligned into a common ecological representation, rather than treated as independent or conflicting sources of information.

This alignment is achieved through a shared encoder trained jointly on observations from all gears, which learns latent features representing underlying ecological gradients common across datasets (Bengio et al., 2013; Pan & Yang, 2010; Reichstein et al., 2019). By optimizing predictions across gears simultaneously, the model emphasizes generalizable ecological patterns while implicitly accounting for gear-specific observation differences, improving cross-domain generalization (Ganin et al., 2016; Pichler et al., 2023). In this way, gear-related differences are treated as variation in observation processes rather than differences in ecological reality.

The transfer learning framework further addresses data imbalance, allowing species-level predictions to leverage broader ecological patterns learned from trophic groups (Pan & Yang, 2010; Weiss et al., 2016). Such approaches are increasingly recognized as critical for ecological modeling in data-limited and heterogeneous systems (Christin et al., 2019; Pichler et al., 2023).

## 5. Conclusion

This study shows that nekton use of seagrass habitats is shaped by both habitat structure and environmental conditions, and that these effects vary across locations. Many previously inconsistent findings in seagrass–nekton relationships can be explained by differences in environmental context and sampling methods, rather than weak ecological relationships.

By combining data collected with different sampling gears, our approach identified consistent ecological patterns across sites while accounting for differences in how species are observed. In practical terms, this means that information from multiple monitoring programs can be used together to better understand habitat use, rather than being analyzed separately or requiring strict standardization.

The results also highlight that environmental gradients (e.g., temperature, salinity, water clarity) set the broad conditions for species distributions, while seagrass structure (e.g., shoots, canopy, biomass) influences habitat use at finer scales. Because these relationships vary across regions, habitat–species relationships are inherently context dependent, and cannot be assumed to be the same across sites.

These findings have direct implications for coastal monitoring, conservation, and restoration. First, they emphasize the value of consistent and coordinated sampling across sites, which improves the ability to detect general ecological patterns (Underwood et al., 2000).

Second, they demonstrate that habitat–species relationships are context dependent, suggesting that restoration strategies must account for environmental gradients and connectivity rather than relying solely on local habitat characteristics (Unsworth et al., 2015; Sievers et al., 2024; Preston et al., 2025).

More broadly, this work provides a framework for integrating diverse ecological datasets to support data-driven decision-making. Such approaches are increasingly important for guiding conservation and restoration under ongoing environmental change driven by climate and land-use pressures (Waycott et al., 2009; Orth et al., 2006).

